# An essential membrane protein modulates the proteolysis of LpxC to control lipopolysaccharide synthesis in *Escherichia coli*

**DOI:** 10.1101/2020.02.10.942425

**Authors:** Elayne M. Fivenson, Thomas G. Bernhardt

## Abstract

Gram-negative bacteria are surrounded by a complex cell envelope that includes two membranes. The outer membrane prevents many drugs from entering these cells and is thus a major determinant of their intrinsic antibiotic resistance. This barrier function is imparted by the asymmetric architecture of the membrane with lipopolysaccharide (LPS) in the outer leaflet and phospholipids in the inner leaflet. The LPS and phospholipid synthesis pathways share a common intermediate. Proper membrane biogenesis therefore requires that the flux through each pathway be balanced. In *Escherichia coli*, a major control point in establishing this balance is the committed step of LPS synthesis mediated by LpxC. Levels of this enzyme are controlled through its degradation by the inner membrane protease FtsH and its presumed adaptor protein LapB(YciM). How turnover of LpxC is controlled has remained unclear for many years. Here, we demonstrate that the essential protein of unknown function YejM(PbgA), which we have renamed ClxD (<u>c</u>ontrol of <u>L</u>p<u>x</u>C <u>d</u>egradation), participates in this regulatory pathway. Suppressors of ClxD essentiality were identified in *lpxC* and *lapB*, and LpxC overproduction was shown to be sufficient to allow survival of Δ*clxD* mutants. Furthermore, the half-life of LpxC was shown to be reduced in cells lacking ClxD, and genetic and physical interactions between LapB and ClxD were detected. Taken together, our results are consistent with a model in which ClxD directly modulates LpxC turnover by FtsH-LapB to regulate LPS synthesis and maintain membrane homeostasis.

**SIGNIFICANCE:** The outer membrane is a major determinant of the intrinsic antibiotic resistance of Gram-negative bacteria. It is composed of both lipopolysaccharide (LPS) and phospholipid, and the synthesis of these lipid species must be balanced for the membrane to maintain its barrier function in blocking drug entry. In this report, we identify an essential protein of unknown function as a key new factor in maintaining LPS/phospholipid balance in the model bacterium *Escherichia coli*. Our results provide novel insight into how this organism and most likely other Gram-negative bacteria maintain membrane homeostasis and their intrinsic resistance to antibiotics.

## INTRODUCTION

The bacterial cell envelope is essential for maintaining cell shape, withstanding mechanical stress, resisting osmotic pressure, and is a bacterium’s first line of defense against antibiotics, bacteriophages, and immune cells (1, 2). In Gram-negative bacteria, the cell envelope includes a symmetric inner membrane (IM) composed of phospholipids and an asymmetric outer membrane (OM) consisting of phospholipids in the inner leaflet and lipopolysaccharide (LPS) in the outer leaflet (2). The cell wall is located in the periplasmic space between the IM and OM and is made from a crosslinked heteropolymer called peptidoglycan (PG). Biogenesis of the cell envelope is tightly coordinated with cellular growth and division (3), with all three layers having to expand and divide each cell cycle without compromising their structural integrity.

The phospholipid (PL) and LPS membrane components are synthesized in the cytosol and within the inner leaflet of the IM. They must be transported across the IM and through the periplasm to build the OM (for reviews, see (1, 4–6)). LPS consists of three covalently attached groups: a glycolipid called Lipid A, a core oligosaccharide, and a longer, variable O-antigen polysaccharide chain. O-antigen is synthesized separately from the Lipid A-core, but the two components are joined by ligation while anchored in the outer leaflet of the IM. Mature LPS molecules are then transported from the IM to the OM by the Lpt system, which forms a protein bridge connecting the IM and OM (7). The mechanism by which PLs are transported to the OM remains unclear (8).

R-3-hydroxymyristoyl-Acyl Carrier Protein (ACP) serves as the acyl donor for the synthesis of both PL and LPS (**Fig. 1A**). In the PL synthesis pathway, it is a substrate of (3R)-hydroxymyristoyl-(acyl-carrier-protein) dehydratase (FabZ) (9). It is also utilized by LpxA (UDP-N-acetylglucosamine acyltransferase) in the first step of Lipid A synthesis (10). However, this reaction is reversible (11); the committed step for Lipid A production is catalyzed by the second enzyme, UDP-3-O-acyl-N-acetylglucosamine deacetylase (LpxC) (12). Balanced synthesis of LPS and PL is required to prevent loss of membrane integrity and cell death (13). In *Escherichia coli*, this balance is in part maintained by the inner membrane-localized, ATP-dependent, zinc-metalloprotease FtsH (14). In conjunction with its presumed adapter protein LapB(YciM) (15, 16), FtsH degrades LpxC to regulate the flux of lipid precursors through the LPS pathway. However, how LpxC proteolysis is modulated in response to disruptions in LPS or PL synthesis to maintain homeostasis has remained unclear for some time.

**Figure 1.**
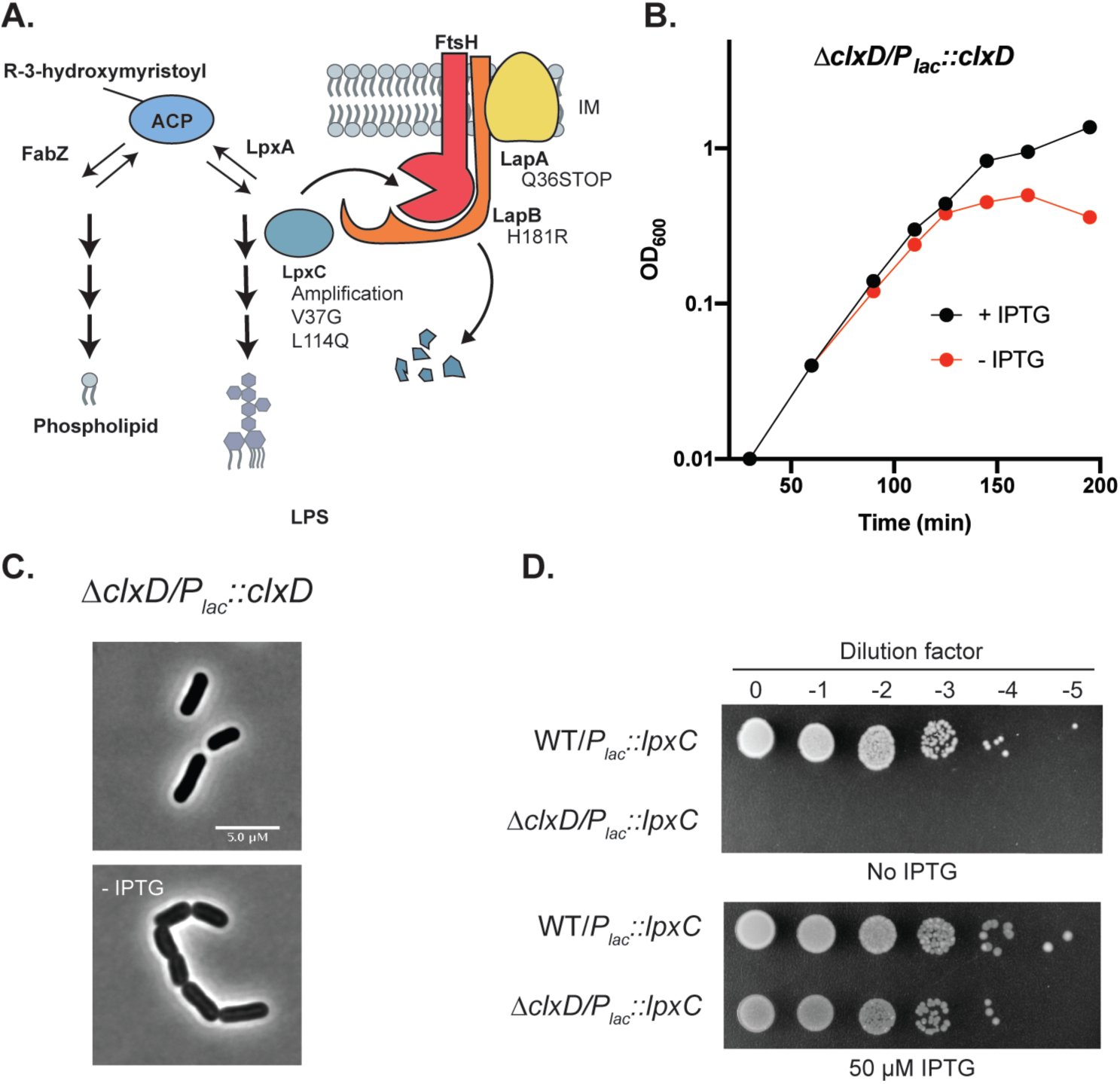
The depletion of ClxD leads to cell chaining and lysis. **A.** Shown is the branched synthesis pathway leading from R-3-hydroxymyistoyl-ACP to either phospholipid or LPS. LpxC is turned over by FtsH. LapB is thought to serve as an adapter in the process. LapA is also shown, but its role in LpxC degradation is not clear. The identity of variants that suppress ClxD essentiality are indicated in the diagram. **B.** Growth curve following the depletion of ClxD. Cells of strain EMF27 [Δ*clxD/P_lac_::clxD*] were grown without (red) or with (black) 50 µM IPTG to induce *clxD* expression and growth was monitored by following culture OD_600_. **C.** Micrographs of cells at the 125 minute time point of the growth curve shown in **B**. Scale bar = 5 µm **D.** Serial dilutions of wildtype and Δ*clxD* cells harboring a *P_lac_::lpxC* plasmid were plated in the absence and presence of 50 µM IPTG as indicated.

Here, we identify the essential membrane protein of unknown function YejM (PbgA) as an inhibitor of LpxC turnover by FtsH-LapB in *E. coli*. We therefore have renamed the protein ClxD for <u>c</u>ontrol of <u>L</u>p<u>x</u>C <u>d</u>egradation. Genetic suppressors of ClxD essentiality first pointed us towards its potential role in maintaining sufficient LpxC levels for LPS synthesis. Subsequent analysis indicated that LpxC half-life is reduced in the absence of ClxD and that ClxD interacts genetically and physically with LapB. Our results are therefore consistent with a model in which ClxD interferes with LpxC proteolysis through its interaction with LapB. We propose that this activity of ClxD is likely to be homeostatic and responsive to perturbations in the balance between LPS and PL synthesis.

## RESULTS

### Rationale

In a genetic selection for suppressors of cell morphogenesis defects in *E. coli*, we identified mutations in *yejM* (renamed *clxD*). These suppressors will be reported as part of a separate study, but they prompted us to further investigate ClxD (YejM, PbgA) function, especially because it has remained one of the few essential genes in *E. coli* without a well-characterized activity. ClxD is an IM protein with a five-transmembrane domain N-terminus that is essential for growth and a non-essential C-terminal periplasmic domain (17, 18). Nonsense mutations in *clxD* leading to the truncation of the periplasmic domain have previously been found to cause phenotypes consistent with defects in OM assembly (17–20), including reduced LPS/PL ratio, vancomycin sensitivity, temperature sensitivity, and leakage of periplasmic proteins. ClxD shares structural similarities to LtaS, the enzyme that synthesizes lipoteichoic acids in many Gram-positive bacteria (21). Like LtaS, ClxD has a hydrophobic binding pocket in the periplasmic domain that is important for protein function (22). However, the crystal structure of this domain of ClxD revealed that it lacks residues that are important for LtaS catalytic activity, indicating that is unlikely to have a similar enzymatic function (22). Although previous studies have implicated ClxD in the transport of cardiolipin to the OM in *Salmonella typhimurium* (23) and *Shigella flexneri* (24), a more recent study has suggested it plays a broader yet ill-defined role in envelope assembly (25). We therefore thought that further study of the function of this essential protein was warranted.

### Overproduction of LpxC suppresses the essentiality of ClxD

To begin investigating the essential function of ClxD, we examined the terminal morphological phenotype induced upon its depletion. The native *clxD* gene was deleted in a strain harboring 6 a plasmid that expressed *clxD* from an IPTG-inducible *lac* promoter (P*_lac_*). In the presence of inducer, these cells grew and divided normally (**Fig. 1B**). However, as observed previously (17, 18, 23), ClxD depletion in the absence of IPTG led to slowed growth followed by lysis of the culture (**Fig. 1B**). Prior to lysing, the ClxD-depleted cells formed cell chains (**Fig. 1C**), indicating a failure to complete cell division. This morphology is reminiscent of cells defective in envelope biogenesis, including mutants defective in LPS assembly (26–28) and cells treated with the LpxC inhibitor CHIR-090 (**SI Appendix, Fig. S1**).

We next turned to suppressor analysis to identify mutations that bypass the essentiality of ClxD as a means to understand its function. To this end, the *clxD* gene was cloned under control of the lactose promoter (P*_lac_*) in a plasmid backbone designed by the de Boer laboratory to select for suppressors of *ftsN* essentiality (29). The vector encodes a temperature-sensitive replication protein (*repA*^ts^) and the restriction endonuclease I-SceI under control of a temperature-sensitive lambda repressor (*cI857*), which is paired with an I-SceI cut site in the vector backbone. To select for suppressors of ClxD essentiality, a Δ*clxD* strain harboring the *clxD* suicide vector was plated at the non-permissive temperature (37°C) where the plasmid will cease to replicate and be cleaved by expressed I-SceI. Twelve surviving colonies were isolated and their mutations were mapped by whole-genome sequencing (**Fig. 1A, SI Appendix,** **Table S1**). Five of the suppressors contained genomic amplifications in a region containing *lpxC*. An additional two suppressors had missense mutations in *lpxC*, including one encoding a V37G substitution that has been demonstrated to lead to increased LpxC abundance in *Klebsiella pneumoniae,* suggesting increased stability (30). Three of the suppressors had mutations either in *lapB*, its neighbor *lapA,* or in the region upstream of the *lapAB* operon. Overall, 10/12 suppressors had mutations predicted to increase cellular LpxC levels.

The suppressor analysis suggested that an LpxC deficiency underpinned the lethality of a Δ*clxD* mutation. To investigate this possibility further, *lpxC* was cloned under P*_lac_* control on a multicopy plasmid. A strain harboring this vector was used as a recipient in a transduction using P1 phage grown on a Δ*clxD::Kan^R^* donor strain. Kanamycin-resistant transductants were readily obtained on agar containing IPTG to induce *lpxC* expression from the plasmid, and these strains were found to be inducer-dependent for growth (**Fig. 1D)**. Notably, the only previously reported suppressor of a Δ*clxD* mutation was a multicopy vector encoding holo-[acyl carrier protein] synthase 2 (AcpT) (17–19), and catalytic activity of the synthase was found not to be required for suppression (17). Given our results, we wondered whether overproduction of AcpT also acts by promoting the accumulation of LpxC. Cells overexpressing *acpT* from a plasmid were indeed found to have elevated LpxC levels (**SI Appendix, Fig. S2**), suggesting that elevated AcpT may overwhelm the FtsH proteolytic machinery and therefore allow for LpxC accumulation. Taken together, our results thus far indicate that LpxC overproduction renders the normally essential ClxD protein dispensable for growth.

### LpxC is aberrantly degraded in the absence of ClxD

To further investigate the connection between ClxD and LpxC, we investigated the effect of ClxD inactivation on the steady-state levels of LpxC. Immunoblotting indicated that LpxC levels were unaffected in wildtype cells with or without moderate induction of *clxD* expression from a plasmid (**Fig. 2A**). However, depletion of ClxD in cells deleted for the native gene led to a dramatic reduction in LpxC levels to the point where it was barely detectable (**Fig. 2A**). Furthermore, increased expression of *clxD* from a plasmid with higher concentrations of inducer led to a striking increase in LpxC accumulation in otherwise wildtype cells (**Fig. 2B**). Based on these results, we hypothesized that LpxC is aberrantly degraded in the absence of ClxD. To test this possibility, we took advantage of our ability to delete *clxD* in the presence of a plasmid overproducing *lpxC*. Wildtype and Δ*clxD* cells expressing *lpxC* from the plasmid were treated with spectinomycin to block translation, and the fate of previously translated LpxC was monitored by immunoblotting. In wildtype cells, the LpxC concentration decayed and then plateaued at a lower steady-state level (**Fig. 2C**). By contrast, LpxC levels decayed rapidly in the Δ*clxD* cells to undetectable levels without an observable plateau. Based on these results, we conclude that ClxD is required to prevent excessive turnover of LpxC.

**Figure 2.**
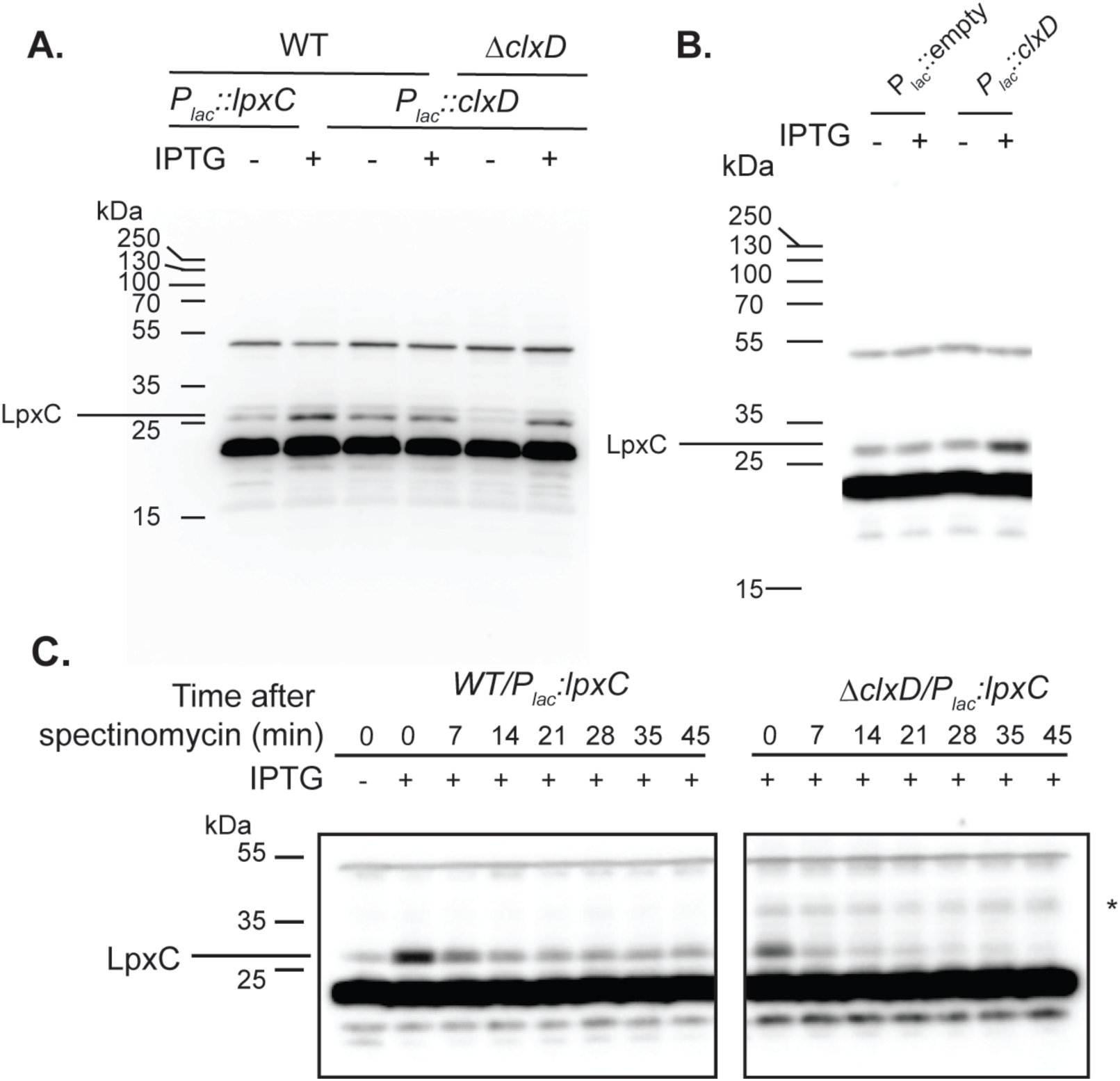
Changes in *clxD* expression affect LpxC accumulation. **A.** Immunoblot using anti-LpxC primary antibody. Lanes 1-4: Extracts from wildtype cells with a *P_lac_::lpxC* (pPR111) or *P_lac_::clxD* (pEMF17) plasmid grown in the absence (lane 1 and 3) or presence (lane 2 and 4) of 50 µM IPTG. Lanes 5 and 6: Extracts from Δ*clxD* cells with a *P_lac_::clxD* (pEMF17) plasmid grown in the absence (lane 5) or presence (lane 6) of 50 µM IPTG. **B.** Immunoblot detecting LpxC from extracts of wildtype cells with a P*_lac_*::empty (pPR66) or a *P_lac_::artificialRBS_clxD* (pEMF33) plasmid grown in the absence (lanes 1 and 3) or presence (lanes 2 and 4) of 1 mM IPTG. Note that the *clxD* in this plasmid has a strong artificial ribosome-binding site relative to the construct used in **A**.**C.** Wildtype and Δ*clxD* cells harboring a *P_lac_::lpxC* plasmid (pPR111) were grown in the presence of 50 µM IPTG at 30°C in minimal medium to an OD_600_ of 0.5. Spectinomycin (300 µg/mL) was then added, and samples were taken at the indicated time points for the preparation of extracts and the detection of LpxC by immunoblotting. The (*) denotes a nonspecific band that appeared sporadically in blots of the Δ*clxD* samples. Blots shown are representative of three independent experiments.

### ClxD interacts genetically and physically with LapB

Our results thus far suggested a model in which ClxD promotes LpxC accumulation by protecting it from the FtsH-LapB proteolytic system. We therefore investigated whether ClxD interacts with any of the components of this system using our recently developed POLAR (PopZ-linked Apical Recruitment) two-hybrid assay (31). The POLAR assay takes advantage of the ability of the PopZ protein from *Caulobacter crescentus* to spontaneously forms foci at the poles of *E. coli* cells. Therefore, protein-protein interactions can be assessed by fusing a “bait” protein a PopZ-interaction domain called H3H4 along with GFP and then monitoring whether it can recruit a “prey” protein of interest fused to mScarlet to the cell pole. Using ClxD as a bait, we found that it was able to recruit a LapB prey to the pole (**Fig. 3B**), but not an FtsH or a LapA prey fusion (**Fig. 3C-D**). The recruitment of LapB prey to the pole was specific for ClxD as it was not recruited by a transmembrane control bait (**Fig. 3A**). However, we noticed that cells expressing LapB-prey constructs formed chains and appeared to lyse when they were paired with the control bait (**SI appendix, Fig. S3**) but not with the ClxD bait. Therefore, for the control in **panel 3A**, we used a control bait construct in which *clxD* was silently expressed downstream of the control bait sequence, which eliminated the adverse morphological effects observed upon production of the LapB-prey fusion. We conclude that ClxD interacts with the LapB component of the FtsH-LapB proteolytic system.

**Figure 3.**
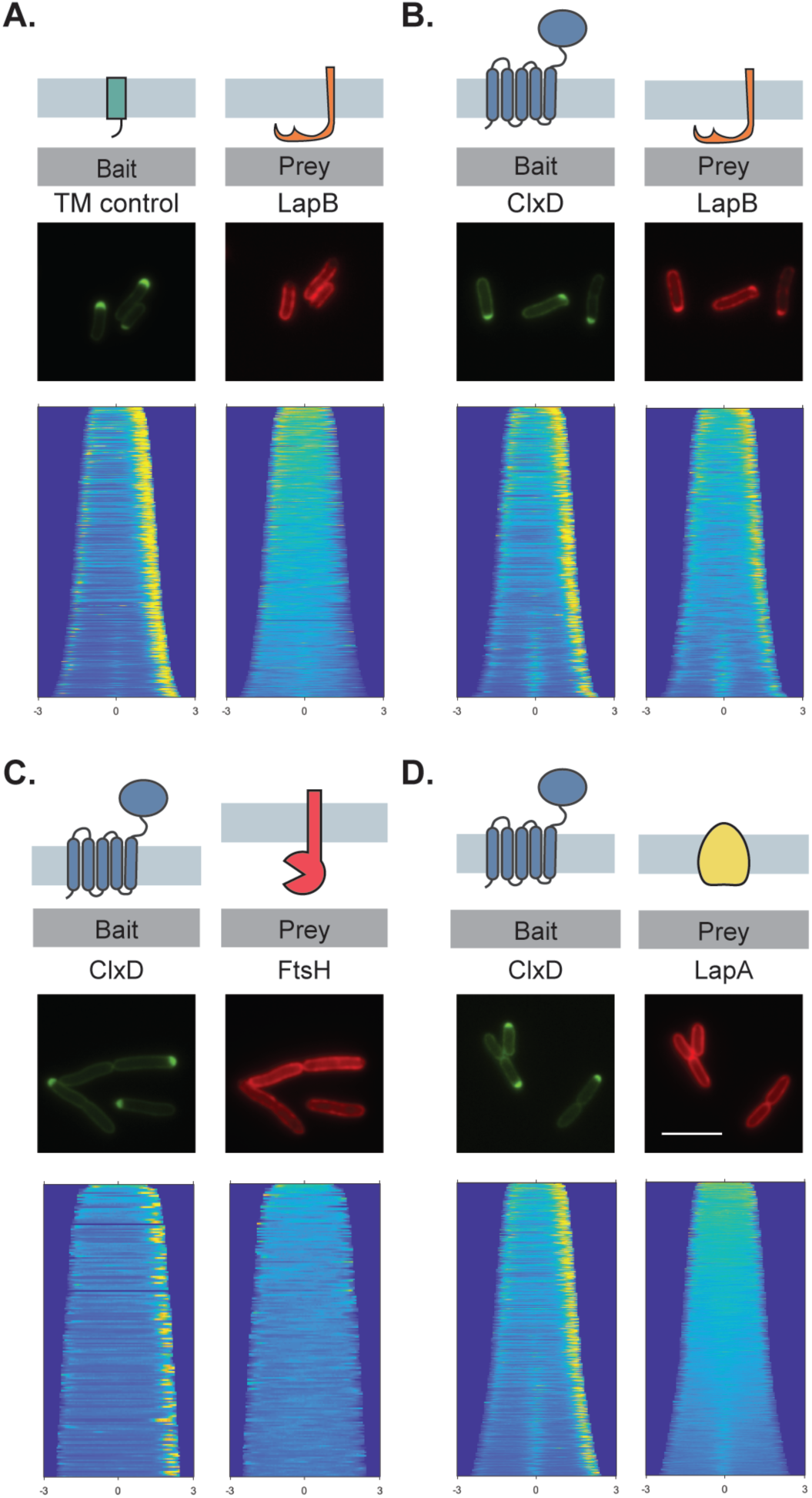
ClxD interacts with LapB. The POLAR assay was used to assess protein-protein interactions (see text for details). Shown are representative micrographs of cells expressing the indicated bait and prey proteins. Cells were transformed with plasmids producing the control bait, which consists of a single transmembrane domain derived from residues 2-55 of *Pseudomonas aeruginosa* PBP1b fused to PopZ-H3H4-GFP (pEMF55) (**A**) or PopZ-H3H4-GFP-ClxD (pEMF35) (**B-D**). Note that the control bait construct also expresses unlabeled *clxD* in order to reduce LapB toxicity. Prey constructs produce C-terminal mScarlet fusions to the indicated proteins from plasmid vectors integrated in the chromosome. Scale bar = 5 µm.

The effects of the LapB-prey fusion observed on cell growth during the POLAR analysis suggested that LapB overproduction is toxic and that this toxicity can be overcome by co-expression *clxD*. To investigate this possibility further, we monitored the effect of overproducing untagged LapB on cell growth when either ClxD or a GFP control was simultaneously overproduced from a compatible plasmid. Cell containing the *lapB* expression plasmid grew normally in the absence of inducer regardless of whether they overproduced ClxD or GFP (**Fig. 4A**). When *lapB* expression was induced, plating efficiency was dramatically reduced for cells producing GFP (**Fig. 4A**). However, cells co-producing ClxD were protected and plated at normal efficiency (**Fig. 4A**). To investigate whether the toxicity of LapB and its antagonism by ClxD was related to LpxC turnover, we monitored LpxC abundance in cells overproducing LapB. Cells in which LapB and GFP were co-produced had reduced LpxC levels, whereas cells overexpressing *clxD* had elevated levels of LpxC and no apparent reduction in LpxC levels upon LapB overproduction (**Fig. 4B**). We conclude that ClxD not only interacts with LapB, but that it likely serves as an antagonist of LapB activity to prevent LpxC degradation by the FtsH-LapB proteolytic machinery.

**Figure 4.**
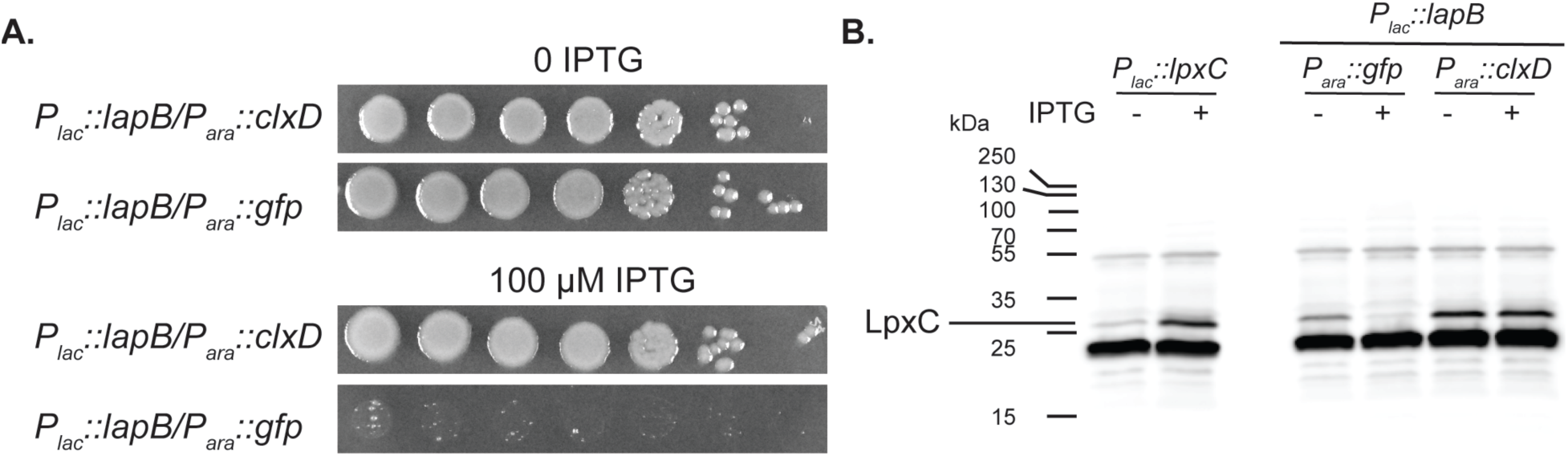
ClxD protects cells from LapB toxicity. **A.** Serial dilutions of wildtype cells with the integrated *lapB* overexpression plasmid pEMF53 [P*_lac_*::*lapB*] and either pEMF57 [P*_ara_*::*gfp*] or pEMF54 [P*_ara_*::*clxD*] were plated on LB agar with 0.2% arabinose additionally supplemented with 100 µM IPTG as indicated. Note that *lapB*-induced toxicity is relieved by *clxD* co-expression. **B**. LpxC immunoblot. Extracts from WT cells harboring a *P_lac_::lpxC* plasmid (pPR111) grown with (Lane 2) or without (lane 1) IPTG were used as a marker for the LpxC band. LpxC was also detected in extracts of MG1655(attHKEMF53) [WT(*P_lac_::lapB*)] cells harboring either pEMF57 [P*_ara_*::*gfp*] (lanes 4 and 5) or pEMF54 [P*_ara_*::*clxD*] (lanes 6 and 7) grown in LB 0.2% arabinose medium without (lanes 4 and 6) or with 100 µM IPTG (Lanes 5 and 7) to induce *lapB* expression.

## DISCUSSION

The function of ClxD (YejM, PbgA) has remained unclear for some time, and it has remained one of the few essential genes of unknown function in *E. coli*. With the exception of a multi-copy suppressor selection (17, 18), prior genetic studies primarily examined the phenotype of mutants encoding a truncated ClxD protein lacking the non-essential C-terminal periplasmic domain (17-20, 23, 25). Cells producing these ClxD-ΔC variants displayed a range of OM permeability barrier defects, suggesting an important yet ill-defined role for ClxD in envelope biogenesis.

A more specific role was assigned to the ClxD homolog from *S. typhimurium*, where it was identified as a factor required for OM barrier function in cells activated for the two-component system PhoPQ (23). This regulatory system induces changes in the OM that help protect *S. typhimurium* from the assaults they encounter in the phagosome of host cells during infection (32). One of the observed changes in OM composition during PhoPQ induction was an increase in the content of the phospholipid cardiolipin (CL) (23, 33). This increase in CL levels in the OM was not observed in PhoPQ-activated cells producing a ClxD-ΔC variant. This observation combined with the detection of CL-binding by the ClxD periplasmic domain in vitro led to the proposal that the protein functions as a transporter that shuttles CL from the IM to the OM (23). In a recent follow-up study (25), mice were infected with *S. typhimurium* producing a ClxD-ΔC variant, and following growth in the host, suppressors that restored OM integrity to the ClxD-ΔC cells were isolated in *lpxC*, *lapB*, and *ftsH* (*25*). It was also found that deletion of the C-terminal domain of ClxD resulted in changes in the LPS and phospholipid composition of *S. typhimurium* that could be at least partially rescued by the suppressors(25). As a result of this analysis, a variety of functions was proposed for ClxD, including a general role in LPS assembly. It was also proposed that the periplasmic domain of ClxD somehow facilitates lipid trafficking during stress and that the transmembrane region performs an undefined essential activity involving phospholipids (25).

A simplified picture of ClxD function emerges from our study of a complete deletion allele of *clxD* in *E. coli*. Similar to the host-induced suppressors of the *S. typhimurium* ClxD-ΔC defect, we identified suppressors of ClxD essentiality in the *lpxC* and *lapB* genes. We further showed that overexpression of *lpxC* was sufficient to allow complete deletion of *clxD*. Subsequent analysis demonstrated that levels of LpxC were strongly dependent on ClxD. Inactivation of ClxD led to increased degradation of LpxC and reduced levels of the enzyme, whereas overproduction of ClxD promoted the hyper-accumulation of LpxC. We then identified an interaction between ClxD and LapB and discovered that ClxD prevents excessive LpxC degradation upon LapB overproduction. Overall, our results are consistent with a model in which ClxD opposes LapB function to inhibit LpxC degradation by the FtsH protease (**Fig. 5**). We therefore suspect that most of the OM defects observed previously for cells producing ClxD-ΔC variants ultimately stem from perturbations to LpxC levels and that a lipid transport function for the protein is unlikely.

**Figure 5.**
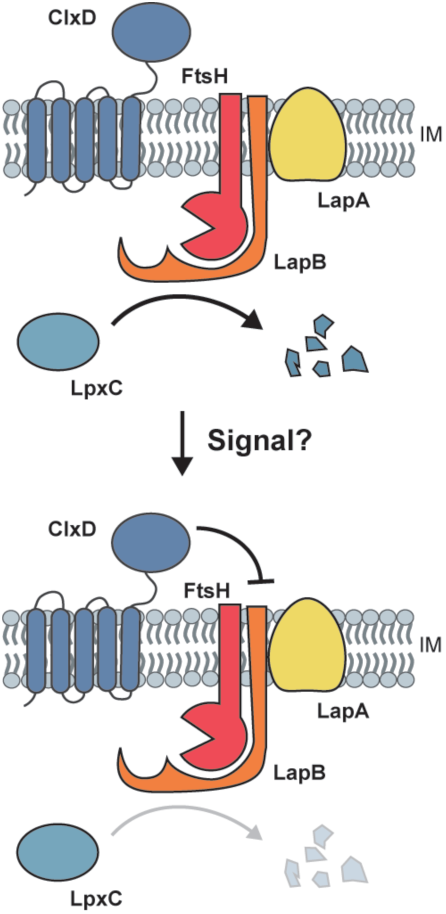
Model for the modulation of LpxC turnover by ClxD. LpxC is degraded by FtsH in a LapB-dependent manner. The role of LapA is unclear. We propose that in response to some signal, potentially the buildup of a lipid molecule in the inner membrane, ClxD blocks LpxC degradation by FtsH-LapB. This inhibition is most likely mediated by an interaction between ClxD and LapB, and may be used to help balance LPS and phospholipid synthesis in response to perturbations/fluctuations.

The ability of ClxD to antagonize LpxC degradation makes it an attractive candidate for a factor that modulates the stability of LpxC in response to disruptions in LPS and phospholipid synthesis homeostasis. It has been known for some time that inhibition of LpxC or overexpression of *fabZ*, both of which presumably shift flux of lipid precursors away from the LPS synthesis pathway, leads to the stabilization of LpxC (9, 30, 34, 35). Recently, it has also been shown that LpxC turnover is reduced in cells with elevated activity of phospholipase A (PldA), which cleaves phospholipids in the outer leaflet of the OM, another marker for defects in LPS and phospholipid synthesis homeostasis (36). In each of these cases, the precise molecular signal(s) that modulate LpxC degradation by FtsH-LapB remains to be determined. However, lipids like acyl-CoA, LPS, phospholipids, or intermediates in the synthesis of these molecules are likely candidates (34, 36). Notably, ClxD is related to enzymes like LtaS and EptA that use phospholipid substrates to either polymerize lipoteichoic acids or to modify Lipid A with a lipid head-group, respectively. Even though ClxD lacks amino acids predicted to be required for enzymatic activity, it is conceivable that it retains the ability to bind LPS, phospholipids, or both, and that such binding events modulate its ability to interfere with LpxC degradation by FtsH-LapB. Although further studies are required to test these possibilities, the identification of ClxD as an essential component in the pathway controlling LpxC levels represents an important step towards a mechanistic understanding of how gram-negative bacteria balance phospholipid and LPS synthesis to properly assemble to OM layer that defines them.

## METHODS AND MATERIALS

### Bacterial Strains and Growth Conditions

All strains used and generated here are derivatives of MG1655. Strains were cultured in LB media (1% tryptone, 0.5% yeast extract, 0.5% NaCl) or minimal M9 media (37) supplemented with 0.2% casamino acids and 0.2% glucose, arabinose, or maltose as indicated. Antibiotic concentrations are as follows: 25 µg/mL chloramphenicol (Cam), 25 µg/mL kanamycin (Kan), 5 µg/mL tetracycline (Tet), 50 µg/mL spectinomycin (Spec), unless otherwise indicated. Strains EMF27 [**Δ***clxD::kan^R^/P_lac_::clxD*] and EMF30 [**Δ***clxD::kan^R^/P_lac_::lpxC*] were maintained on medium supplemented with 50 µM IPTG unless otherwise stated. All strains, plasmids, and primers used in this study are listed in **SI Appendix** **Table S2**, **SI Appendix** **Table S3**, and **SI Appendix** **Table S4**, respectively. Methods used to construct the Δ*clxD* strains and expression constructs used in this study are detailed in the **SI Appendix**.

### Selection of suppressors of ClxD essentiality

The suppressor strain plasmid (pEMF20) was cloned via Gibson assembly in strain JLB45, which expresses *cI*857, in order to prevent zygotic induction of *I-SceI*. The plasmid was then transformed into MG1655 Chung competent cells. The Δ*clxD::kan^R^* allele was transduced into MG1655/pEMF20 via P1 transduction and confirmed via PCR (see above) generating strain EMF37. Suppressors of Δ*clxD* were selected by growing EMF37 cells at 37°C on LB plates (including 1% SDS, as indicated). An overnight culture of EMF37 was prepared at the permissive condition (30°C, LB + Kan + Spec +100 µM IPTG). For suppressors EMF40-44, the EMF37 overnight culture was serially diluted and plated on LB plates and incubated at 37°C. For suppressors EMF45-51, the overnight culture of EMF37 was back diluted 1:50 and grown under permissive conditions until OD_600_ = 0.42. Cells were then pelleted, washed, and resuspended in LB and allowed to grow for an additional hour at 37°C (non-permissive conditions). Cells were then plated on LB and incubated overnight at 37°C. The loss of plasmid pEMF20 was confirmed by screening for spectinomycin sensitivity. Overnight cultures of the suppressor strains were prepared, and 5 ml of each culture was pelleted and stored at −20°C. gDNA was isolated from each pellet using the Wizard Genomic DNA Purification Kit (Promega) and further purified using the Genomic DNA Clean & Concentrate kit (Zymo Research). Whole genome sequencing was performed as described previously (38) with some modifications (39) (Nextera DNA Sample Preparation kit). The concentration of the DNA in the samples was determined using the Qubit dsDNA HS Assay Kit and the sizes of the products following tagmentation were determined using a High Sensitivity D1000 screen tape run on an Agilent 4200 TapeStation. The sequencing was carried out using a MiSeq Reagent Kit v3 (Illumina). The data was analyzed using the CLC Genomics Workbench software (Qiagen).

### Immunoblotting

Cell pellets were collected and resuspended in water and 2x Laemmli sample buffer [100 mM Tris-HCL, pH 6.8; 2% SDS, 0.1% Bromophenol Blue, 20% glycerol] at a 1:1 ratio. Samples were boiled for 10 minutes and sonicated (Qsonica Tip Sonicator-Amp: 25%, Time: 1 min) 2-3 times. Protein concentration was measuring using the NI (Non-interfering) Protein Assay (With BSA Protein Standard)(G Biosciences cat #786-005) and was normalized using 1x sample buffer. Samples were run on a 15% polyacrylamide gel and transferred to a PVDF membrane. The membrane was then rinsed in Phosphate-Buffered Saline + 0.1% tween (PBS-T) [10% 10x PBS buffer PBS-T, pH 7.4(Sigma Aldrich); 0.1% Tween 20) and blocked in 5% milk in PBS-T for 1.5 hrs. The membrane was then incubated in primary antibody solution of 1% milk in PBS-T + rabbit anti-lpxC antibody (generous gift from the Doerrler lab) at a 1:10,000 dilution for approximately 16 hours at 4°C. The membrane was then washed 4x in PBS-T (1x quickly, 3x for 10 min per wash). The membrane was then incubated in secondary antibody solution (anti-rabbit IgG HRP, 1:1000 dilution, Rockland 18-8816-33) in 0.2% milk in PBS-T for 2 hrs. Following 5 washes with PBS-T, the membrane was developed using SuperSignal West Pico PLUS Chemiluminescent Substrate (Thermo Fisher Scientific cat#34577) and imaged using the c600 Azure Biosystems platform.

### LpxC degradation assay

MG1655/pPR111 [WT/*P_lac_::lpxC*] and EMF30 [Δ*clxD/P_lac_::lpxC*] cells were incubated overnight in 4 mL of LB + Cam with or without 50 µM IPTG, respectively. Overnight cultures were diluted to OD_600_ = 0.025 in 5 mL of LB + Cam with (MG1655/pPR111 and EMF30) or without (MG1655/pPR111) 50 µM IPTG. Cultures were incubated at 37°C with shaking until OD_600_ ∼0.3-0.4. Cell concentrations were then normalized and cultures were then back diluted again at a ratio of 1:50 in 100 mL M9 minimal media + 50 µg/mL chloramphenicol and 50 µM IPTG (except for the MG1655/pPR111 control without IPTG). Cells were grown at 30°C with shaking until OD_600_ = 0.5. Then, 300 µg/mL spectinomycin was added to the cultures to inhibit protein synthesis. Samples were taken at 0, 7, 14, 21, 28, 35, 45 minutes following the addition of spectinomycin and analyzed via immunoblotting for LpxC.

### Monitoring Growth during ClxD depletion

Cultures of TB28/pEMF17 [Δ*lacIZYA::frt*/*P_lac_::clxD*] and EMF27 [Δ*lacIZYA::frt, ΔclxD/P_lac_::clxD*] were grown overnight at 37°C in 4 mL of LB + Cam + 50 µM IPTG. Cultures were then diluted to OD_600_=0.025 in 6 mL of LB + Cam + 50 µM IPTG and grown at 37°C with shaking until OD_600_=0.3. Then, 3 mL of culture was pelleted for 2 min at 5000 rpm, washed 1x in LB + Cam, and resuspended in 3 mL LB + Cam. The concentration of the cultures was then normalized and the samples were diluted at a ratio of 1:50 in 50 LB + Cam +/-50 µM IPTG. Cultures were incubated at 37°C with shaking. Samples were taken every 15-30 minutes as indicated. The culture density was measured and samples of cells were removed for fixation. Growth curve data was then plotted in Graphpad Prism.

### Effect of CHIR-090 on cell morphology

MG1655 [WT] cells were grown overnight in LB at 37°C and back diluted to OD_600_ = 0.05 in 5 mL of LB and grown until OD_600_ = 0.4. Cells were back-diluted again to OD_600_ = 0.1 in 5 mL LB + DMSO or 0.25 µg/mL CHIR-090 (Caymen Chemical Company cat #728865-23-4). Cells were grown for 90 minutes and fixed before being visualized by phase contrast microscopy.

### Phase Contrast Microscopy

For the phase contrast micrographs in **Fig. 1** and **SI Appendix Fig. S1**, cells were fixed in 2.6% in formaldehyde and 0.04% glutaraldehyde at room temperature for 1 hr and stored at 4°C for up to three days. Prior to imaging, samples were immobilized on 2% agarose pads on 1 mm glass slides and # 1.5 coverslips were used. Samples were imaged on a Nikon TE2000 inverted microscope using Nikon Elements Acquisition Software AR 3.2. Cropping and additional adjustments were made using FIJI software.

### POLAR analysis

Cells were prepared for imaging as described previously (31). Briefly, cells from a single colony were grown in LB media supplemented with Tet and Cam for 2 hours at 37°C. Cells were back-diluted in M9 + 0.2% arabinose and 100 µM IPTG and incubated for 2 hours at 37°C to induce expression of bait and prey protein fusions. Cells were immobilized on agarose pads as above for imaging. Micrographs were taken on a Nikon Ti Inverted Microscope with a Plan Apo lambda 100x/1.45 Oil Ph3 DM objective lens, Lumencore SpectraX LED Illumination, Chroma ET Filter Cubes GFP (49002) and mCherry (49008), a Andor Zyla 4.2 Plus sCMOS Camera, and Nikon Elements 4.30 acquisition software. The microscope slide was kept at 30°C using an environmental control chamber. Demographs showing polar localization were generated using a custom MATLAB code as described previously (31). Cells were aligned by length and oriented so that the pole with greater bait intensity is located on the right. The corresponding demograph of the prey signal was generated using the same orientation.

## ACKNOWLEDGEMENTS

We would like to thank all the members of the Bernhardt and Rudner Labs for their thoughtful discussions and advice throughout this project, especially Patty Rohs for her insight and guidance in the early stages of this project. We would also like to acknowledge the consultation, support, and services provided by Paula Montero Llopis and the other members of MicRoN (Microscopy Resources on the North Quad) team at HMS. We thank Bill Doerrler for the generous gift of the anti-LpxC antibody. This work was supported by the National Institutes of Health (AI083365 to T.G.B) and Investigator funds from the Howard Hughes Medical Institute. E.M.F. was supported in part by the T32 Bacteriology PhD Training Program (AI132120-02) awarded to the Harvard Graduate Program in Bacteriology

## Supplemental Material

**Figure S1.**
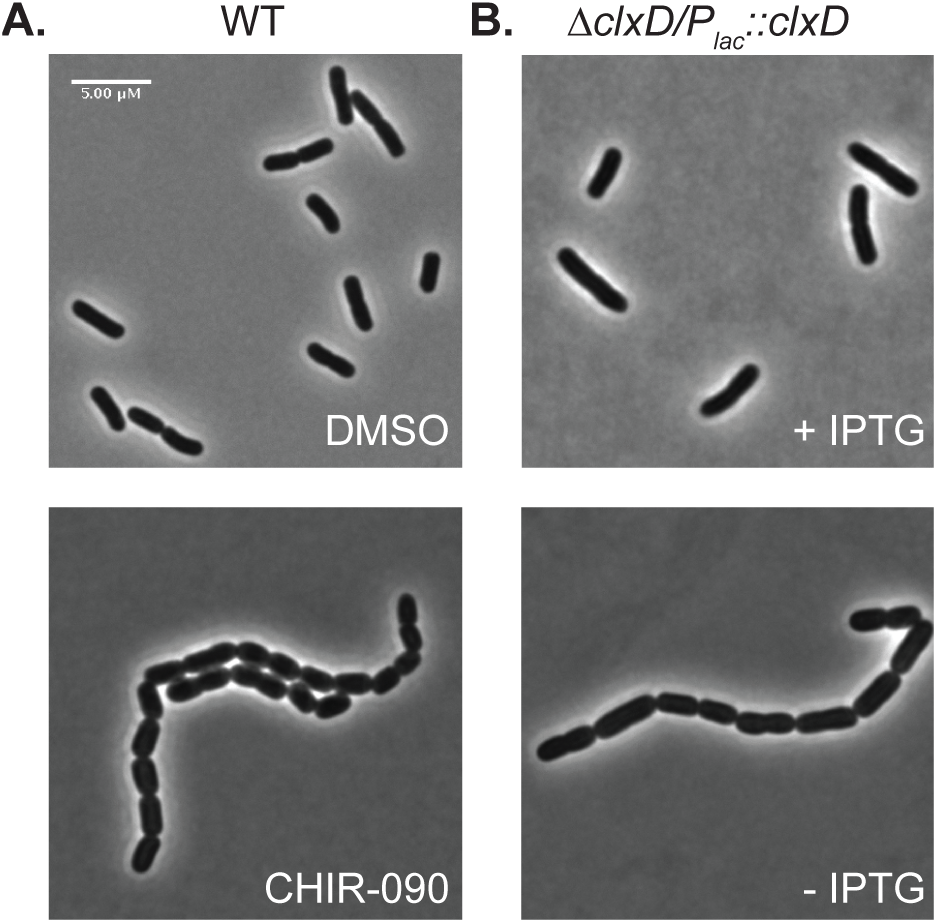
ClxD-depleted cells have a similar morphology to cells treated with the LpxC inhibitor CHIR-090. **A.** Wildtype cells treated with DMSO (top) or 0.25 µg/mL CHIR-090 (bottom) for 2 hrs. **B**. Δ*clxD/P_lac_::clxD* cells 145 minutes after growth in the presence or absence of 50 µM IPTG (top) as indicated. Scale bar = 5 µm.

**Figure S2.**
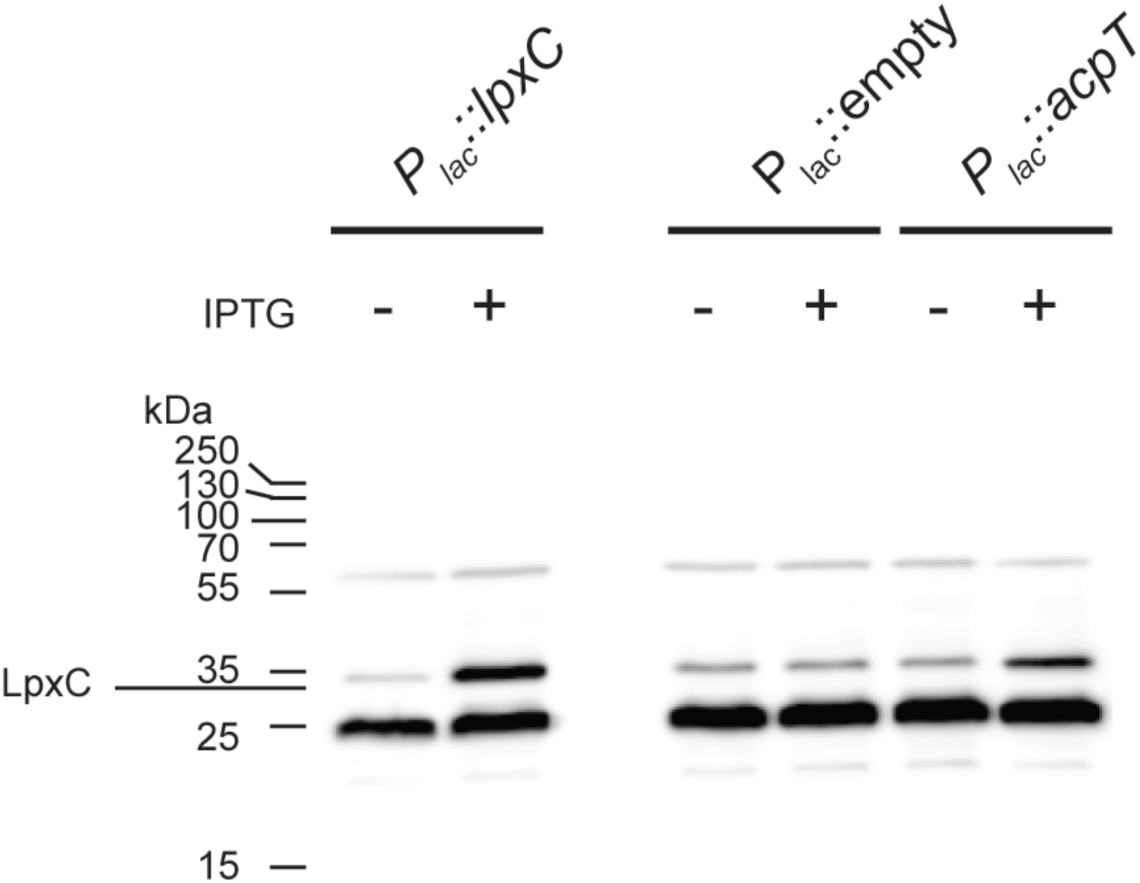
The overexpression of *acpT* increases LpxC levels. Immunoblot detection of LpxC from extracts of wildtype cells harboring plasmids pPR111 [P*_lac_*::*lpxC*], pPR66 [P*_lac_*::*empty*], or pEMF43 [P*_lac_*::*acpT*] grown in the presence or absence of IPTG (50 µM lane 2, or 1 mM for lanes 4 and 6) as indicated.

**Figure S3.**
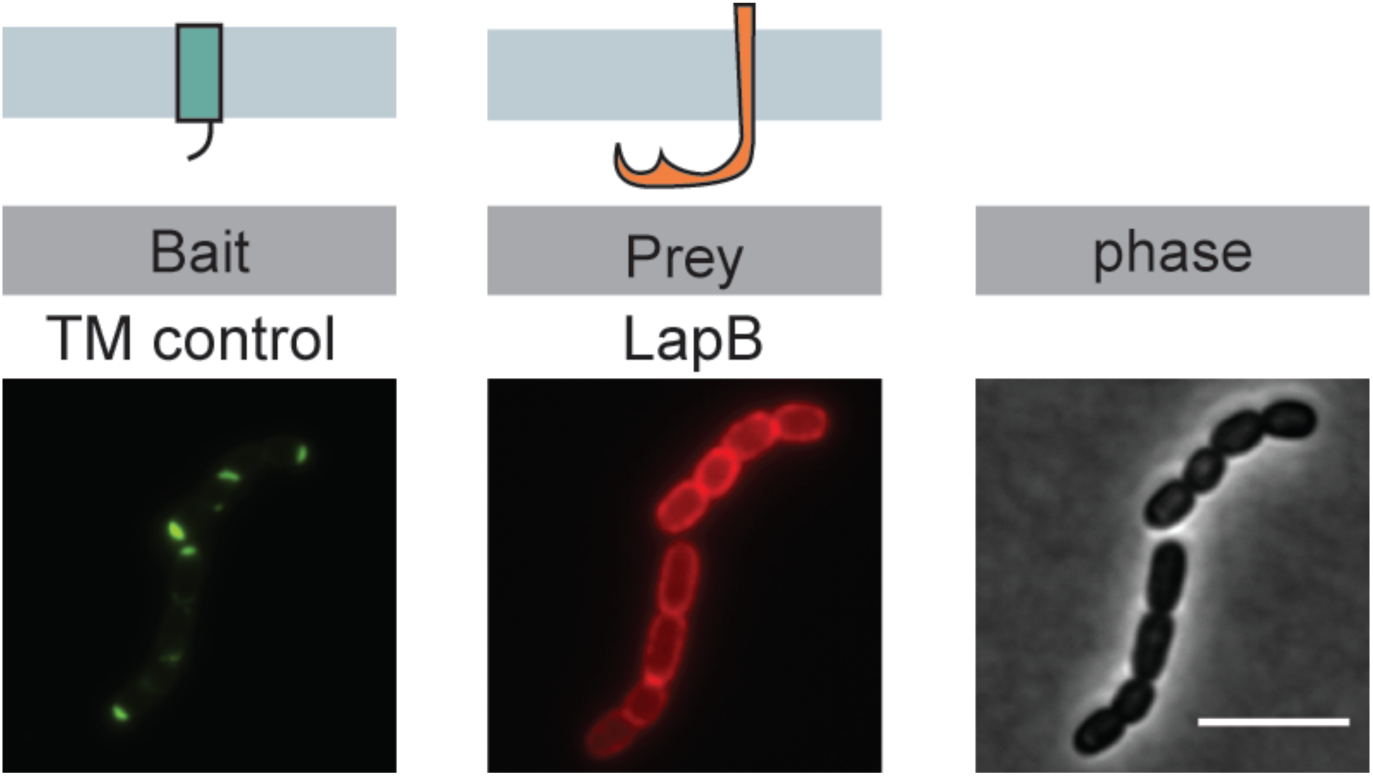
The LapB prey construct does not interact with control bait construct in the POLAR assay. Shown are representative micrographs of cells expressing the indicated bait and prey proteins. Cells were transformed with plasmids expressing the control bait, which consists of a single transmembrane domain derived from residues 2-55 of *Pseudomonas aeruginosa* PBP1b fused to PopZ-H3H4-GFP (pHCL149). The LapB prey construct consists of a C-terminal mScarlet fusion to LapB and was expressed from an integrated plasmid (pHCL147 derivative pEMF36) under the P*_lac_* promoter. The experimental setup is the same as for Figure 3A, except unlabeled ClxD is not expressed from the bait construct in this case. Note that without expressing the unlabeled ClxD, cells begin to form chains, most likely due to the toxic effects of LapB overproduction. Scale bar = 5 µM.

**Table S1.** Δ*clxD* suppressors.

| **Strain** | **Selection condition** | **Mutation** | **Potential causative mutation** |
| --- | --- | --- | --- |
| EMF40 | LB 37°C | Amplification (*yaiO-yjgX*) | Amplification of *lpxC* |
| EMF41 | LB 37°C | *aspC*(A371A) (silent mutation)<br>*lpxC*(V37G) | Point mutant in *lpxC* |
| EMF42 | LB 37°C | Amplifications (*ykfC-rrlE*) and (*hemG-yhcF*) | Amplification of *lpxC* |
| EMF43 | LB 37°C | Amplification (*hokE-yjgX*), A-> T point mutation upstream of *gspH* | Amplification of *lpxC* |
| EMF44 | LB 37°C | Amplifications (*hoke-nmpC*) and (*ykfC-mokC*) | Amplification of *lpxC* |
| EMF45 | LB 37°C 1% SDS | *lapB*(H181R) | *lapB*(H181R) |
| EMF46 | LB 37°C 1% SDS | C-> A point mutant upstream of *lapAB* operon | C-> A point mutant upstream of *lapAB* operon |
| EMF47 | LB 37°C 1% SDS | *lpxC*(L114Q) | *lpxC*(L114Q) |
| EMF48 | LB 37°C 1% SDS | *lapA*(Q36QSTOP) | *lapA*(Q36QSTOP) |
| EMF49 | LB 37°C 1% SDS | Amplification (*yejO-yecF*) | unknown |
| EMF50 | LB 37°C 1% SDS | Amplification (*yejO-yecF*) | unknown |
| EMF51 | LB 37°C 1% SDS | Amplification (*yaiX- yjgX*) | Amplification of *lpxC* |

**Table S2.**
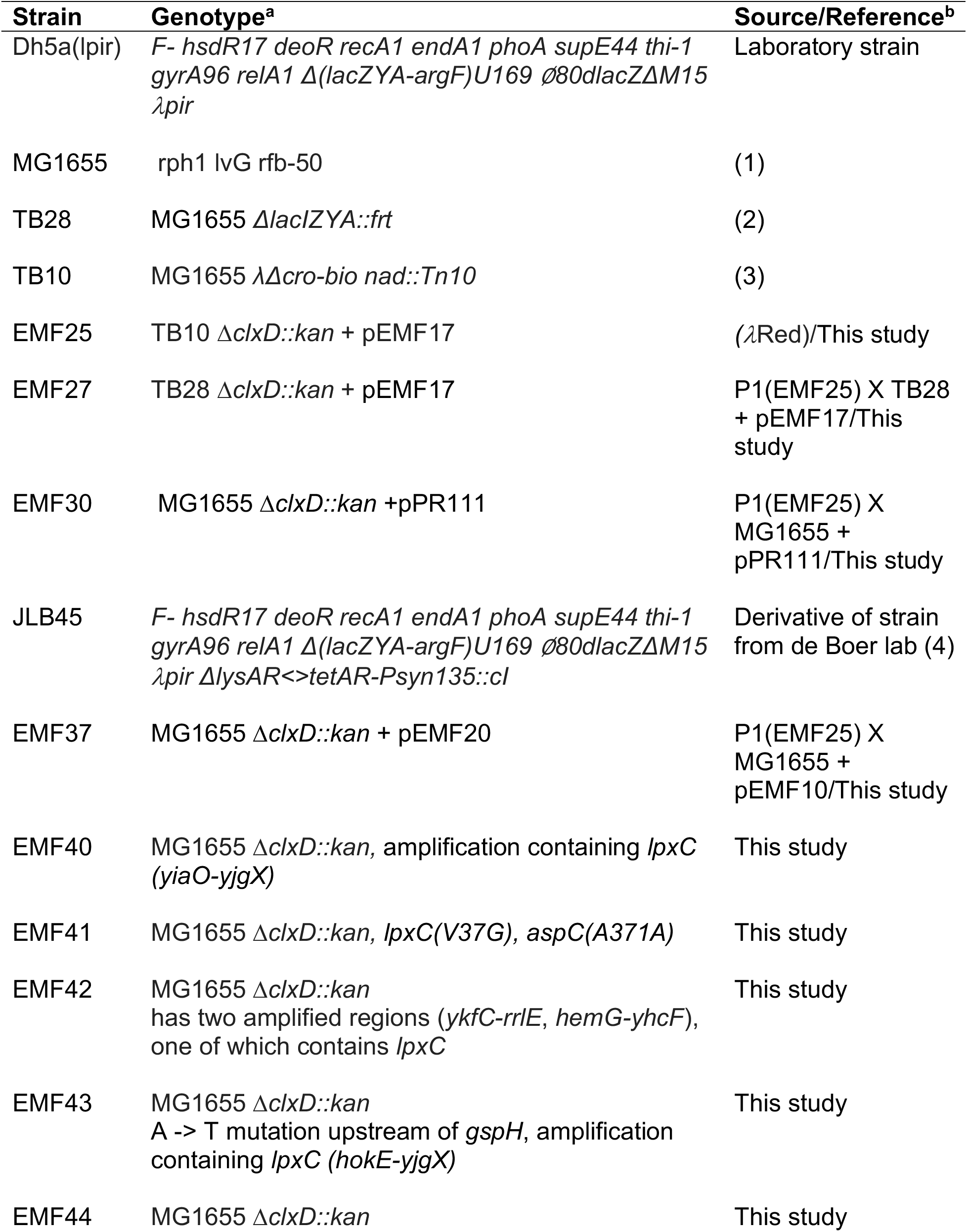
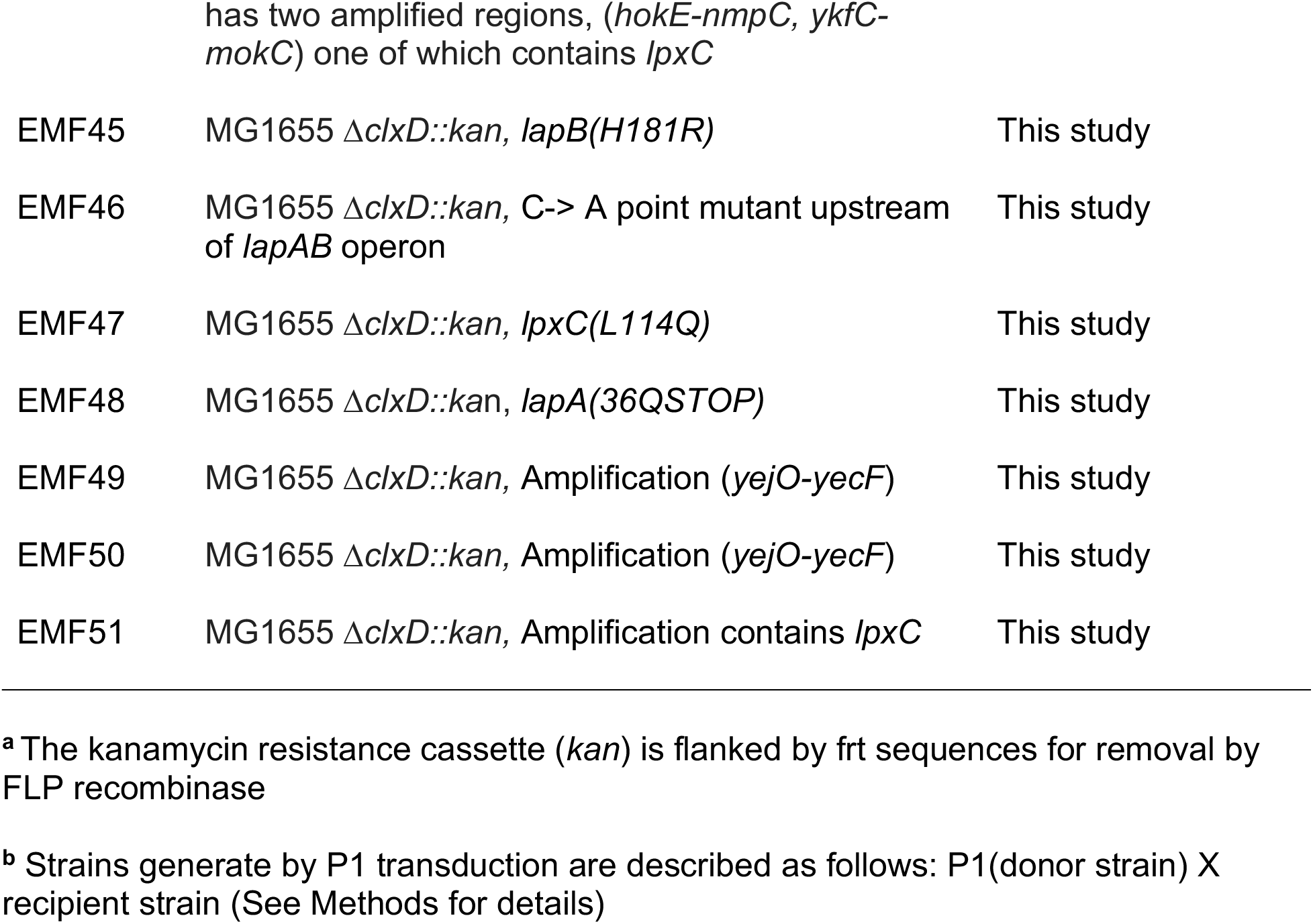
Strains used in this study.

**Table S3.**
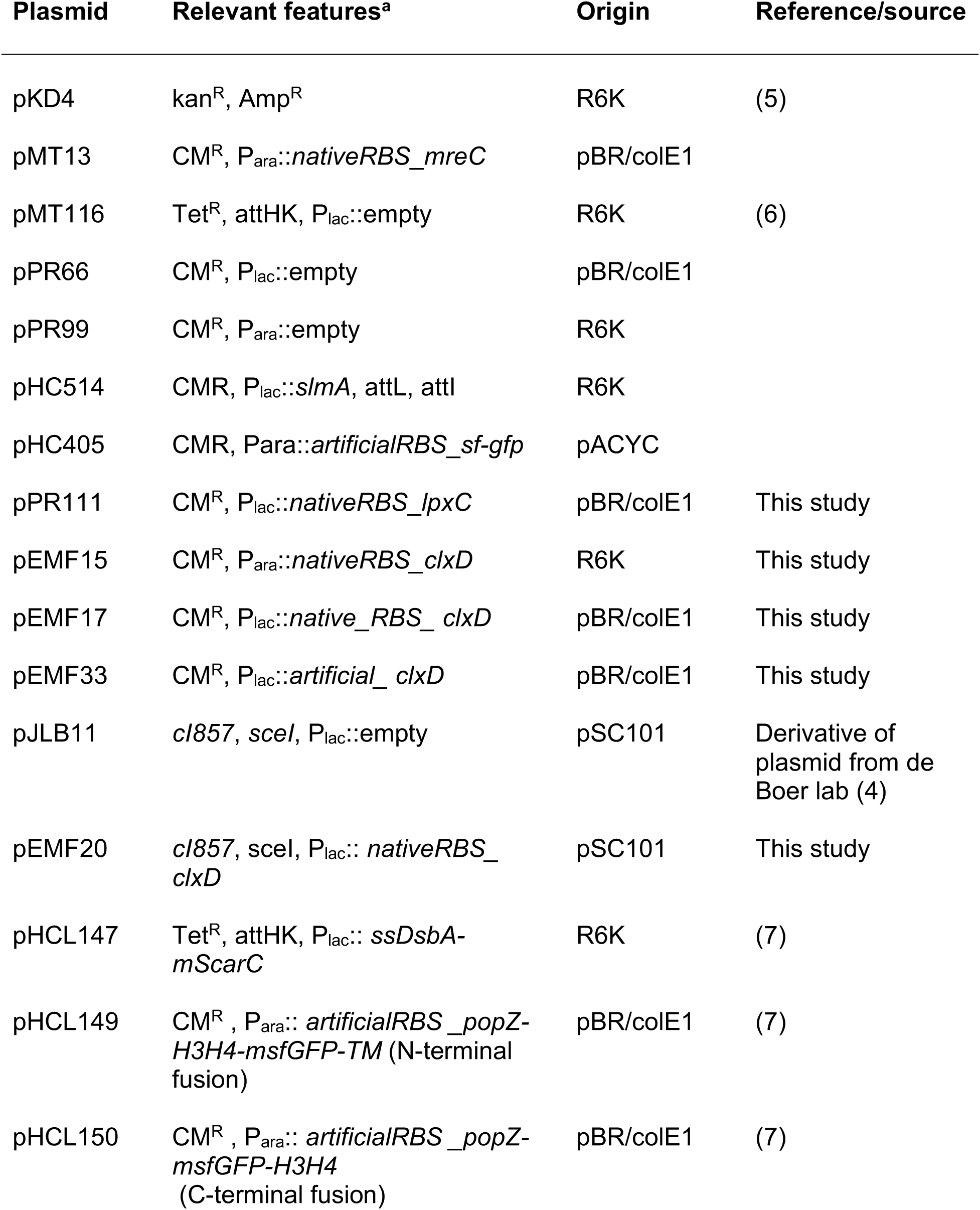
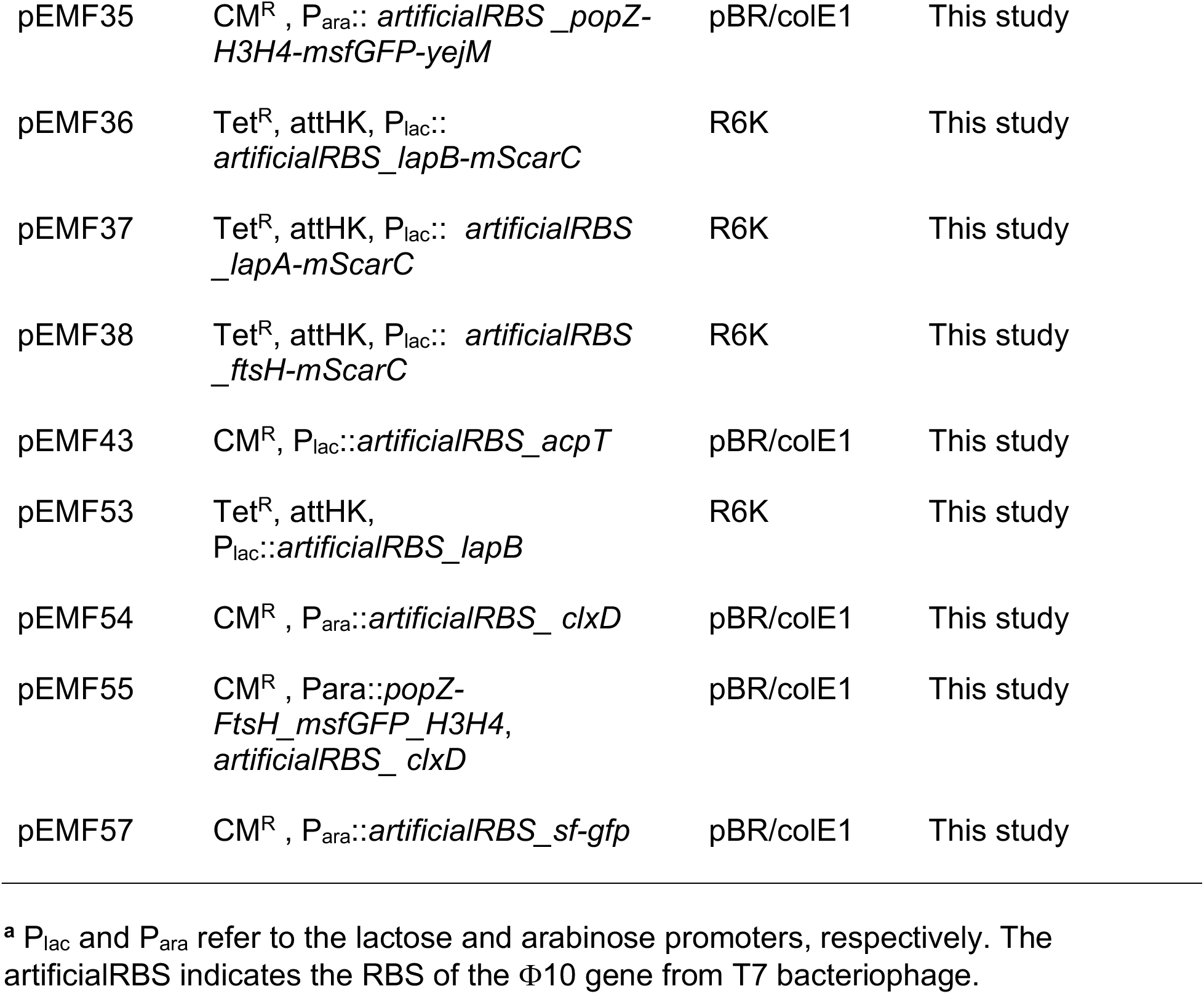
Plasmids used in this study.

**Table S4.**
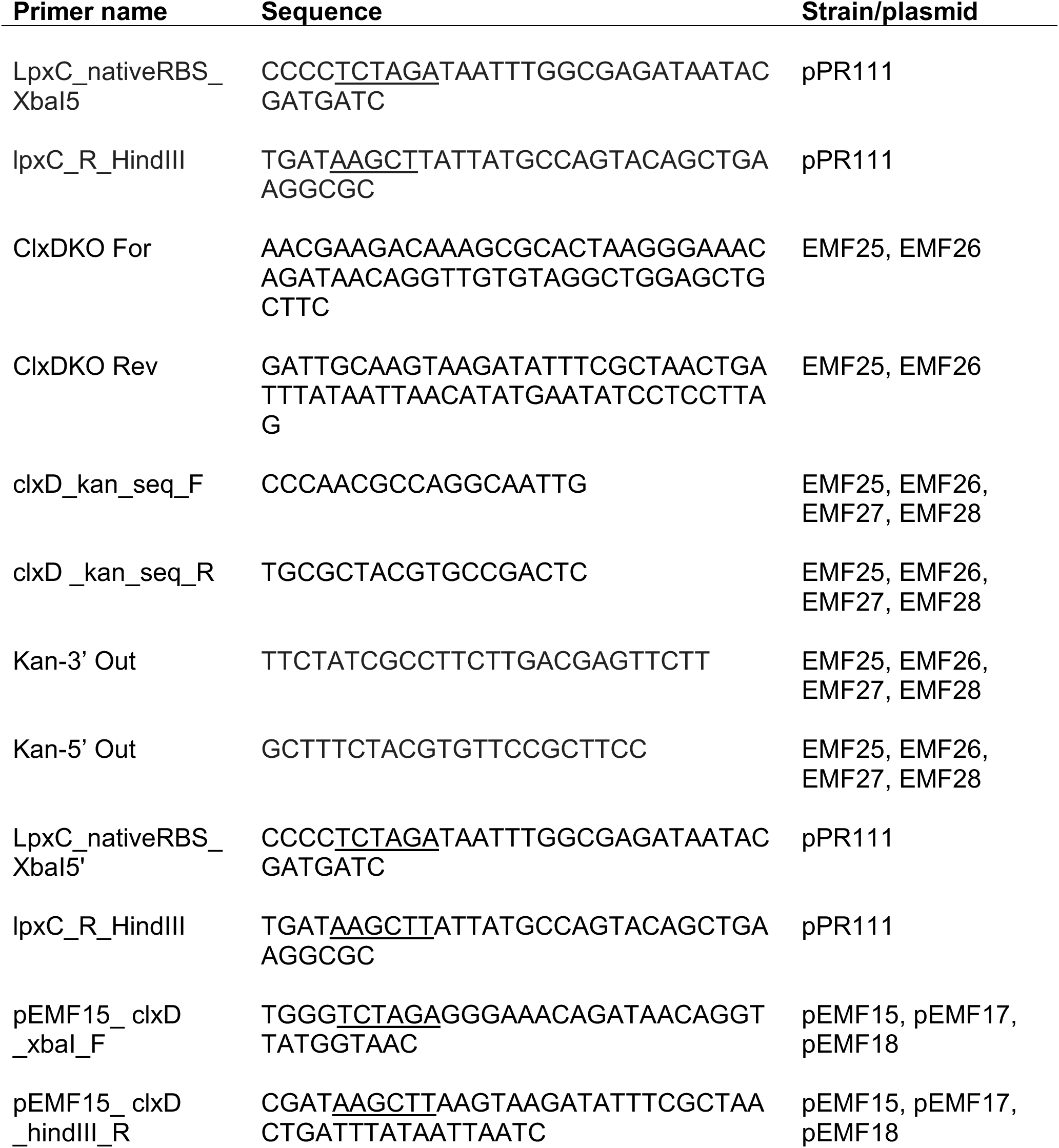
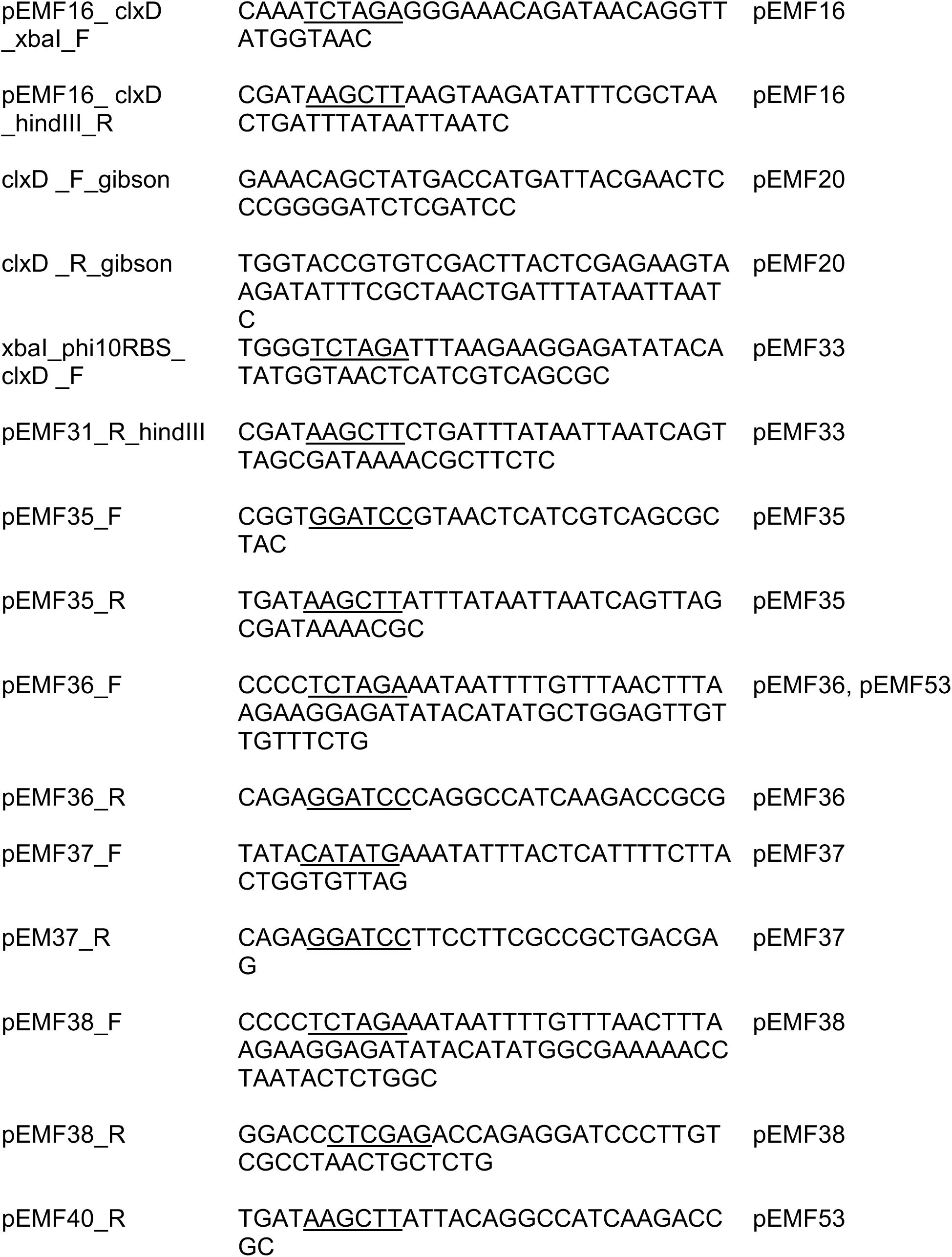
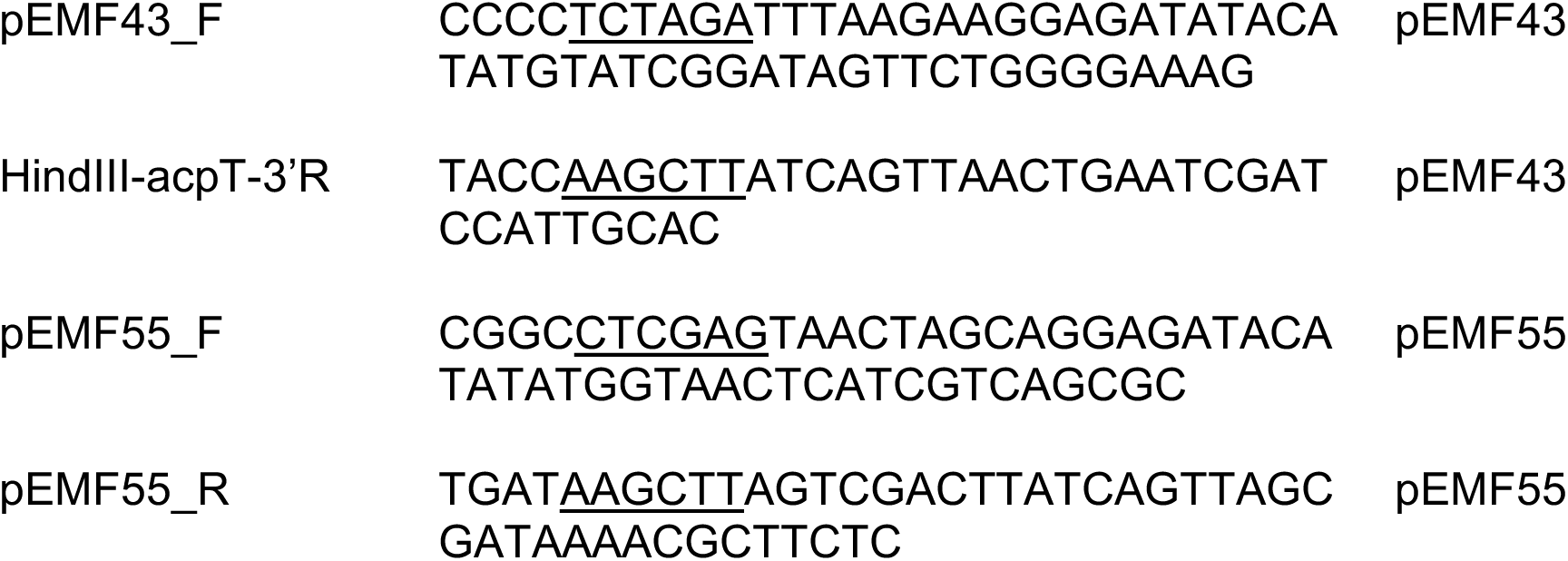
Primers used in this study.

## MATERIALS AND METHODS

### Construction of the ΔclxD strain EMF27 [ΔclxD::kan^R^/P_lac_::clxD]

The *kan^R^* cassette and flanking FRT sites was amplified from pKD4 using primers with homology domains flanking *clxD* coding sequence (forward primer: AACGAAGACAAAGCGCACTAAGGGAAACAGATAACAGGTTGTGTAGGCTGGAGCT GCTTC, reverse primer: GATTGCAAGTAAGATATTTCGCTAACTGATTTATAATTAACATATGAATATCCTCCTTAG. This PCR product was then transformed into TB10 (8) harboring plasmid pEMF17 [P_lac_::*clxD*]. Transformants were selected on LB + Kan + Cm + IPTG (100 µM). The Δ*clxD::kan^R^* allele was confirmed by PCR across the 5’ junction using the primers (clxD_kan_seq_F: CCCAACGCCAGGCAATTG, Kan-5’ Out: GCTTTCTACGTGTTCCGCTTCC) and 3’ junction (clxD_kan_seq_R: TGCGCTACGTGCCGACTC, Kan-3’ Out: TTCTATCGCCTTCTTGACGAGTTCTT) and for lack of growth in the absence of inducer. The Δ*clxD::kan^R^* allele was then transduced from EMF25 into TB28/pEMF17 to generate strain EMF27. The genotype was confirmed by PCR and lack of growth in the absence of inducer.

### Construction of the ΔclxD strain EMF30 [ΔclxD::kan^R^/P_lac_::lpxC]

The Δ*clxD::kan^R^* allele was transduced into MG1655/pPR111 [WT/P*_lac_*::*lpxC*]. Transductants were selected on LB + Cam + Kan + 50 µM IPTG. The transduction was confirmed via PCR (see above).

### Molecular biology

The polymerase chain reaction (PCR) was conducted using Q5 High fidelity polymerase (New England Biolabs) or GoTaq green master mix (Promega) following manufacturer’s protocol. PCR products were purified using the PCR clean up kit from Qiagen or CWBiosciences. Plasmids were isolated using the Miniprep Kit from Qiagen or the plasmid purification kit from CWBiosciences. All plasmids were sequence verified by the DNA Resource Core of Dana-Farber/Harvard Cancer Center.

### Plasmid construction

#### pPR111

LpxC was amplified from gDNA using primers LpxC_nativeRBS_XbaI5’ (CCCC<u>TCTAGA</u>TAATTTGGCGAGATAATACGATGATC) and lpxC_R_HindIII (TGAT<u>AAGCTT</u>ATTATGCCAGTACAGCTGAAGGCGC). The resulting insert (xbaI_nativeRBS_lpxC_hindII) was cloned into vector pPR66 using restriction enzymes xbaI and hindIII.

#### pEMF15

ClxD was amplified from gDNA using primers pEMF15_clxD_xbaI_F (TGGG<u>TCTAGA</u>GGGAAACAGATAACAGGTTATGGTAAC) and pEMF15_clxD_hindIII_R (CGAT<u>AAGCTT</u>AAGTAAGATATTTCGCTAACTGATTTATAATTAATC). The resulting insert (xbaI_nativeRBS_*clxD*_hindII) was cloned into vector pPR99 using restriction enzymes xbaI and hindIII.

#### pEMF17

pEMF15 was digested with restriction enzymes xbaI and hindIII-HF and the xbaI_nativeRBS_*clxD*_hindIII insert was gel-extracted (see Materials and Methods) and ligated into digested pPR66.

#### pEMF43

acpT was amplified from gDNA using primers pEMF43_F (CCCC<u>TCTAGA</u>TTTAAGAAGGAGATATACATATGTATCGGATAGTTCTGGGGAAAG) and HindIII-acpT-3’R (TACC<u>AAGCTT</u>ATCAGTTAACTGAATCGATCCATTGCAC). The insert (xbaI_artificialRBS_acpT_hindIII) was ligated into pPR66 using using restriction enzymes xbaI and hindIII.

#### pEMF20

*clxD* was amplified from pEMF17 using primers clxD_F_gibson (GAAACAGCTATGACCATGATTACGAACTCCCGGGGATCTCGATCC) and clxD_R_gibson (TGGTACCGTGTCGACTTACTCGAGAAGTAAGATATTTCGCTAACTGATTTATAATTA ATC) and inserted in pJLB11, which was digested with restriction enzymes smaI and xhoI, via isothermal assembly. This plasmid was constructed in strain JLB45 which expresses the *cI*857, in order to prevent zygotic induction.

#### pEMF33

*clxD* was amplified from pEMF17 using primers xbaI_phi10RBS_clxD_F (TGGG<u>TCTAGA</u>TTTAAGAAGGAGATATACATATGGTAACTCATCGTCAGCGC) and pEMF31_R_hindIII (CGAT<u>AAGCTT</u>CTGATTTATAATTAATCAGTTAGCGATAAAACGCTTCTC). The resulting insert (xbaI_artificialRBS_clxD_hindII) was cloned into vector pPR66 using restriction enzymes xbaI and hindIII.

#### pEMF35

*clxD* was amplified from pEMF17 using primers pEMF35_F (CGGT<u>GGATCC</u>GTAACTCATCGTCAGCGCTAC) and pEMF35_R (TGAT<u>AAGCTT</u>ATTTATAATTAATCAGTTAGCGATAAAACGC). The resulting product (BamHI_*clxD*_hindIII) was cloned into vector pHCL149(7) using restriction enzymes BamHI and HindIII-HF.

#### pEMF36

*lapB* was amplified from gDNA using primers pEMF36_F (CCCC<u>TCTAGA</u>AATAATTTTGTTTAACTTTAAGAAGGAGATATACATATGCTGGAGTT GTTGTTTCTG) and pEMF36_R(CAGA<u>GGATCC</u>CAGGCCATCAAGACCGCG). The resulting PCR product (xbaI_artificialRBS_lapB_BamHI) was cloned into pHCL147(7) using restriction enzymes xbaI and BamHI. The ligation was the transformed into TB28/pTB102 Chung competent cells. The insert was then amplified and sequenced. Correct integration was confirmed via PCR (9). The integrated plasmid was then transduced into MG1655. The resulting strain was confirmed via PCR.

#### pEMF37

*lapA* was amplified from gDNA using primers pEMF37_F (TATA<u>CATATG</u>AAATATTTACTCATTTTCTTACTGGTGTTAG) and pEM37_R (CAGA<u>GGATCC</u>TTCCTTCGCCGCTGACGAG). The resulting product (ndeI_lapA_bamHI) was inserted into pHCL147(7) using restriction enzymes ndeI and bamHI. The ligation was the transformed into TB28/pTB102 Chung competent cells. The insert was then amplified and sequenced. Correct integration was confirmed via PCR (9). The integrated plasmid was then transduced into MG1655. The resulting strain was confirmed via PCR.

#### pEMF38

FtsH was amplified from gDNA using primers pEMF38_F (CCCC<u>TCTAGA</u>AATAATTTTGTTTAACTTTAAGAAGGAGATATACATATGGCGAAAAA CCTAATACTCTGGC) and pEMF38_R (GGACC<u>CTCGAG</u>ACCAGAGGATCCCTTGTCGCCTAACTGCTCTG). The resulting product (xbaI_artificialRBS_ftsH_xhoI) was inserted into pHCL147(7) using restriction enzymes xbaI and xhoI. The ligation was the transformed into TB28/pTB102 Chung competent cells. The insert was then amplified and sequenced. Correct integration was confirmed via PCR(9). The integrated plasmid was then transduced into MG1655. The resulting strain was confirmed via PCR.

#### pEMF53

artificalRBS_lapB was amplified from pEMF40 using primers pEMF36_F (CCCC<u>TCTAGA</u>AATAATTTTGTTTAACTTTAAGAAGGAGATATACATATGCTGGAGTT GTTGTTTCTG) and pEMF40_R(TGAT<u>AAGCTT</u>ATTACAGGCCATCAAGACCGC). The resulting insert (xbaI_artificalRBS_lapB_hindIII) was digested using restriction enzymes xbaI and hindIII-HF and ligated into vector pMT116. The ligation was the transformed into TB28/pTB102 Chung competent cells. The insert was then amplified and sequenced. Correct integration was confirmed via PCR(9). The integrated plasmid was then transduced into MG1655. The resulting strain was confirmed via PCR.

#### pEMF54

pEMF33 was digested with restriction enzymes xbaI and hindIII-HF. The artificalRBS_clxD insert was ligated into pMT13.

#### pEMF55

ArtificalRBS_*clxD* sequence was amplified from pEMF35 using primers pEMF55_F(CGGC<u>CTCGAG</u>TAACTAGCAGGAGATACATATATGGTAACTCATCGTCA GCGC) and pEMF55_R(TGAT<u>AAGCTT</u>AGTCGACTTATCAGTTAGCGATAAAACGCTTCTC). The resulting insert (xhoI_artificalRBS_*clxD*_hindIII) was digested using restriction enzymes xhoI and hindIII-HF and ligated into vector pHCL149 directly downstream of the Para::popZ-H3H4-msfGFP-TM sequence.

#### pEMF57

pHC405 was digested with restriction enzymes xbaI and hindIII-HF. The xbaI_sf-gfp_hindIII insert was ligated into pMT13.

